# Harmonization of L1CAM Expression Facilitates Axon Outgrowth and Guidance of a Motor Neuron

**DOI:** 10.1101/2020.06.09.143248

**Authors:** Tessa Sherry, Hannah R. Nicholas, Roger Pocock

## Abstract

Brain development requires precise regulation of axon outgrowth, guidance and termination by multiple signaling and adhesion molecules. How the expression of these neurodevelopmental regulators is transcriptionally controlled is poorly understood. The *Caenorhabditis elegans* SMD motor neurons terminate axon outgrowth upon sexual maturity and partially retract their axons during early adulthood. Here we show that C-Terminal Binding Protein-1 (CTBP-1), a transcriptional corepressor, is required for correct SMD axonal development. Loss of CTBP-1 causes multiple defects in SMD axon development: premature outgrowth, defective guidance, delayed termination and absence of retraction. CTBP-1 controls SMD axon development by repressing the expression of SAX-7 – a L1 cell adhesion molecule (L1CAM). CTBP-1-regulated repression is crucial as deregulated SAX-7/L1CAM causes aberrant SMD axons. We found that axonal defects caused by SAX-7/L1CAM misexpression are dependent on a distinct L1CAM, called LAD-2, which itself plays a parallel role in SMD axon guidance. Our results reveal that harmonization of L1CAM expression controls the development and maturation of a single neuron.

## INTRODUCTION

Establishment of neuronal circuits within the brain requires choreographed events, including axon outgrowth, guidance, fasciculation and termination. Once development is complete, maintenance factors promote stable axon morphology and position throughout life (Aurelio et al., 2002; Sasakura et al., 2005), although examples of structural plasticity such as axon pruning and retraction also exist (Bagri et al., 2003; Luo and O’Leary, 2005; Xu and Henkemeyer, 2009). These complex axonal behaviors are directed by intrinsic and extrinsic molecular interactions and signaling pathways that require precise spatial and temporal control (Hutter, 2019; Tessier-Lavigne and Goodman, 1996). Ultimately, integration and harmonization of these signaling pathways enables axons to correctly navigate complex molecular and cellular environments. However, regulatory mechanisms that govern spatial and temporal control of axon guidance signals are poorly understood.

The L1 family of cell adhesion molecules (L1CAMs) are transmembrane proteins, typically composed of 6 immunoglobulin (Ig) domains, 3-5 fibronectin III domains (FnIII) and a short cytoplasmic domain containing an ankyrin binding motif, FERM domain and PDZ domain (Brummendorf et al., 1998). L1CAMs coordinate multiple adhesion and signaling mechanisms in nervous system development, maintenance and function (Brummendorf et al., 1998; Cohen et al., 1998). As a result, mutations in L1 family members result in a wide range of human neurological abnormalities, including CRASH disorder (corpus callosum hypoplasia, retardation, adducted thumbs, spastic paraplegia, and hydrocephalus) (Nagaraj et al., 2014). Vertebrates typically encode four L1 family members (L1, CHL1, Neurofascin and NrCAM), whereas invertebrates contain one or two L1 orthologs (Brummendorf et al., 1998). The nematode *Caenorhabditis elegans* encodes two L1CAMs, LAD-2 and SAX-7, which play multiple autonomous and non-autonomous functions in axo-dendritic development and maintenance (Benard et al., 2012; Chen et al., 2001; Dong et al., 2013; Pocock et al., 2008; Salzberg et al., 2013; Sasakura et al., 2005; Wang et al., 2008). However, the regulatory mechanisms governing L1CAM expression remain largely elusive.

The SMDDs are a bilaterally symmetric pair of cholinergic motor neurons that extend axons to pioneer the *C. elegans* neuropil (nerve ring) during embryogenesis (Rapti et al., 2017). During post-embryonic development, the SMDDs extend posteriorly-directed axons from the head into the dorsal sublateral nerve cord where they terminate in the anterior half of the animal. The SMDDs innervate dorsal muscles to drive head bending and regulation of omega turn amplitude, and are functionally important for exploratory behavior and proprioception (Cook et al., 2019; Gray et al., 2005; Shen et al., 2016; White et al., 1986; Yeon et al., 2018). Here we show that the C-Terminal Binding Protein-1 (CTBP-1) controls SMDD development by regulating SAX-7/L1CAM expression. We provide genetic and molecular evidence that CTBP-1 repression of *sax-7* is required for SMDD guidance but not outgrowth. We further show that the CTBP-1-SAX-7 regulatory relationship controls SMDD development in a temporally-distinct and parallel pathway to the other *C. elegans* L1CAM ortholog LAD-2. Thus, appropriate expression of two L1CAMs is important to control SMDD development and that CTBP-1-dependent transcriptional repression of SAX-7 permits the axon-promoting function of LAD-2. Taken together, we have discovered a mechanism controlling axonal development by harmonizing L1CAM expression.

## RESULTS

### The SMDD neurons undergo phases of axon outgrowth and retraction

The SMDDs are a pair of cholinergic sublateral motor neurons that extend dorsally-directed axons to pioneer the *C. elegans* nerve ring during embryogenesis (Rapti et al., 2017). We surveyed post-embryonic development of the SMDD neurons using a *Pglr-1::GFP* transgene and found that SMDD axon outgrowth is continuous throughout larval development and the first day of adulthood, though not scaled with the increase in worm body length (Figure 1A-C, S1A and Table S1). We found that termination of SMDD axon outgrowth occurs ~170μm from the terminal bulb of the pharynx in 1-day adults (Figure 1C and Table S1). Subsequently, we observed that during days 1-3 of adulthood, the SMDD axons retract by 14μm (~8% of their length), with no further retraction from days 3-5 (Figure 1C, and Table S1). We detected no associated decrease in body length during these adult stages (Figure S1A and Table S1). These observations suggest that the SMDDs undergo autonomous axonal remodeling during postembryonic development and adulthood.

**Figure 1.**
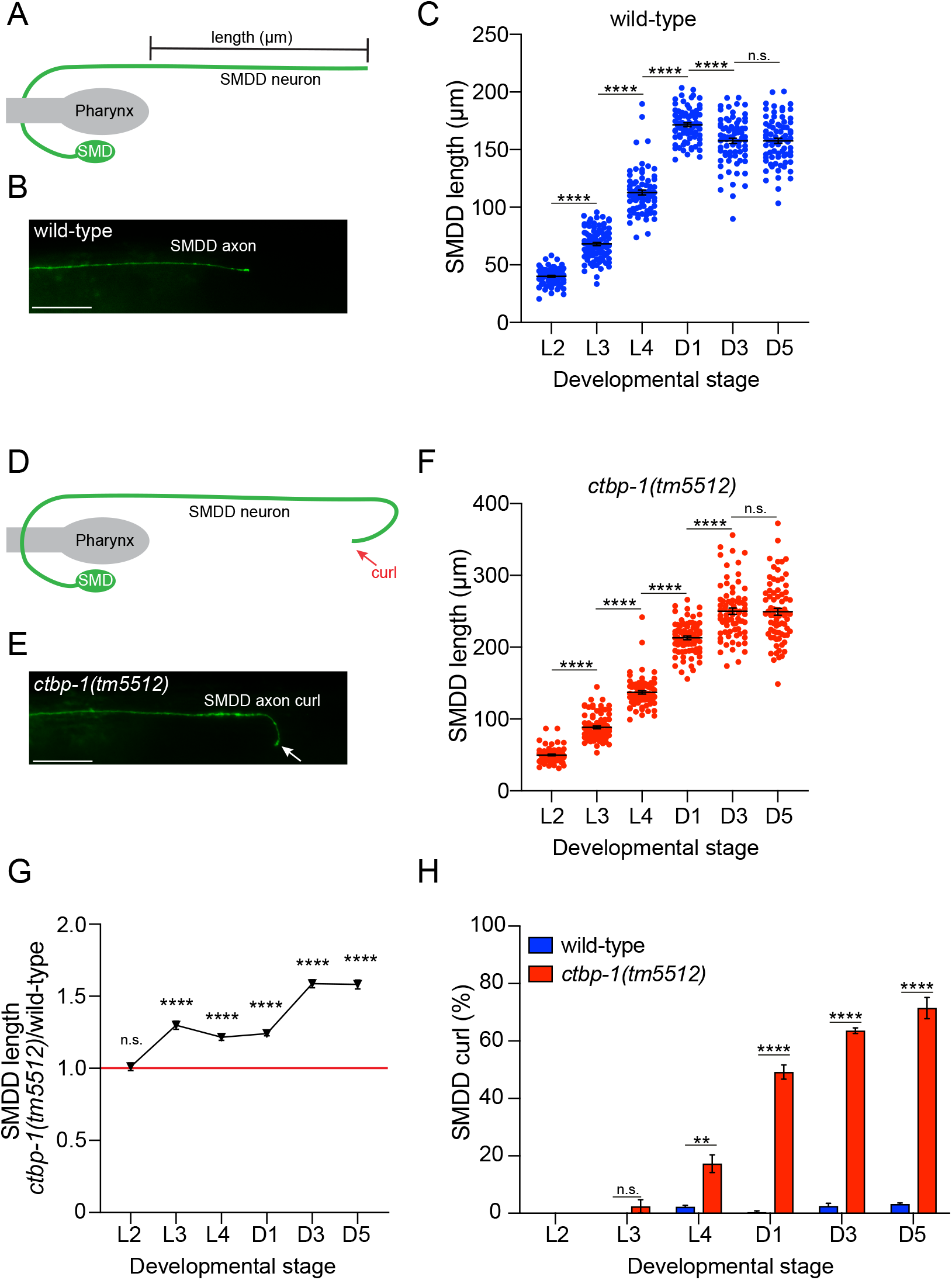
CTBP-1 regulates SMDD outgrowth and retraction. (A) Schematic of SMDD axon morphology in wild-type worms. Bar shows the axon segment measured in C and F. (B) Image of an SMDD axon in a wild-type L4 larva. Scale bar, 20μm. (C) Quantification of SMDD axon length during wild-type post-embryonic development (larval stages L2-L4; adult stages D1, D3 and D5). Data presented as individual axon lengths (points) with mean ± S.E.M (bar). n=75-91 axons, ****p<0.0001, n.s. – not significant (unpaired t-test). (D) Schematic of SMDD axon ‘curl’ defect observed in *ctbp-1(tm5512)* mutant worms. (E) Image of SMDD axon in a *ctbp-1(tm5512)* mutant L4 larva. Arrow indicates the axon curling away from the sublateral nerve cord. Scale bar, 20μm. (F) Quantification of SMDD axon length during *ctbp-1(tm5512)* post-embryonic development (same developmental stages as in C). Data presented as individual axon lengths (points) with mean ± S.E.M (bar). n=73-86 axons, ****p<0.0001, n.s. – not significant (unpaired t-test). (G) Quantification of SMDD axon length of *ctbp-1(tm5512)/wild-type*. Data presented as mean ± S.E.M (bar). n=73-86 axons. ****p<0.0001, n.s. – not significant (unpaired t-test). (H) Quantification of the SMDD axon curl phenotype in wild-type and *ctbp-1(tm5512)* animals during post-embryonic development (same developmental stages as in C). Data presented as mean ± S.E.M (bar) of 3 biological replicates, n>100 axons. **p<0.01, ****p<0.0001, n.s. – not significant (unpaired t-test). In Figure 1, SMDD morphology was visualized by a fluorescent reporter – *rhIs4[Pglr-1::GFP]*.

### CTBP-1a controls SMD axon outgrowth and retraction

We previously reported that the transcriptional corepressor C-Terminal Binding Protein-1 (CTBP-1) is important for SMDD axonal morphology (Reid et al., 2015). Loss of *ctbp-1* causes aberrant SMDD axon guidance where axons turn away (curl) from the dorsal sublateral nerve cord (Figure 1D-E) (Reid et al., 2015). We examined whether CTBP-1 also controls SMDD axon termination/retraction. We found that SMDD axons are longer in *ctbp-1(tm5512)* mutant animals compared to wild-type at all developmental stages from L3 larvae onwards (Figure 1F-G and Tables S1-2). We further show that, unlike wild-type animals, the SMDD axons of *ctbp-1(tm5512)* mutants continue to extend between days 1-3 of adulthood and maintain their length in day 5 adults *(ctbp-1(tm5512)* = 249μm c.f. wild-type = 158μm) Figure 1F and Table S2). These data reveal that in *ctbp-1(tm5512)* mutant animals, the SMDD axons fail to terminate in early adulthood and do not retract as worms age. Increased SMDD axonal length was independent of overall body length, as *ctbp-1a* mutants were either shorter (L2-L4, D3) or had similar body length as wild-type animals (D1 and D5) (Figure S1A and Tables S1-2). Further, the SMDD axons of *ctbp-1(tm5512)* mutants were longer than wild-type irrespective of whether they extended within the sublateral cord or were misguided outside the cord (Figure S1B). As the *ctbp-1(tm5512)* mutant SMDD overgrowth phenotype is already detected in L3 larvae, the curl phenotype, which is only detectable from the L4 stage, may be caused by premature axon outgrowth or a separate mechanism (Figure 1G-H).

The *ctbp-1* locus generates two protein isoforms – CTBP-1a and CTBP-1b. The CTBP-1a isoform contains a sequence-specific THAP (Thanatos-associated protein) DNA binding domain, a PXDLS-binding cleft that potentially coordinates protein-protein interactions, and a nucleotide binding dehydrogenase-like domain (Figure 2A) (Nicholas et al., 2008). CTBP-1b lacks the THAP DNA binding domain and has a specific N-terminal amino acid sequence (Figure 2A). Our data show that the *ctbp-1(tm5512)* mutant, which disrupts the *ctbp-1a*-specific THAP DNA binding domain, exhibits defective SMDD axonal development (Figure 1). We explored whether the THAP DNA binding domain is required for CTBP-1 control of SMDD development by generating a CTBP-1b-specific deletion – *ctbp-1(aus15)* – using CRISPR-Cas9 (Figure 2A). We found that *ctbp-1b(aus15)* mutant SMDD axons terminate and retract normally and do not exhibit the axon curl phenotype (Figure 2B and S1C). Further, animals harboring deletions in both isoforms, *ctbp-1a/b(tm5512aus14)*, exhibit the same axon curl phenotype as the *ctbp-1a(tm5512)* mutant (Figure 2B). These data indicate that CTBP-1a, and not CTBP-1b, is required for SMDD development, suggesting that DNA binding through the THAP domain drives this developmental decision.

**Figure 2.**
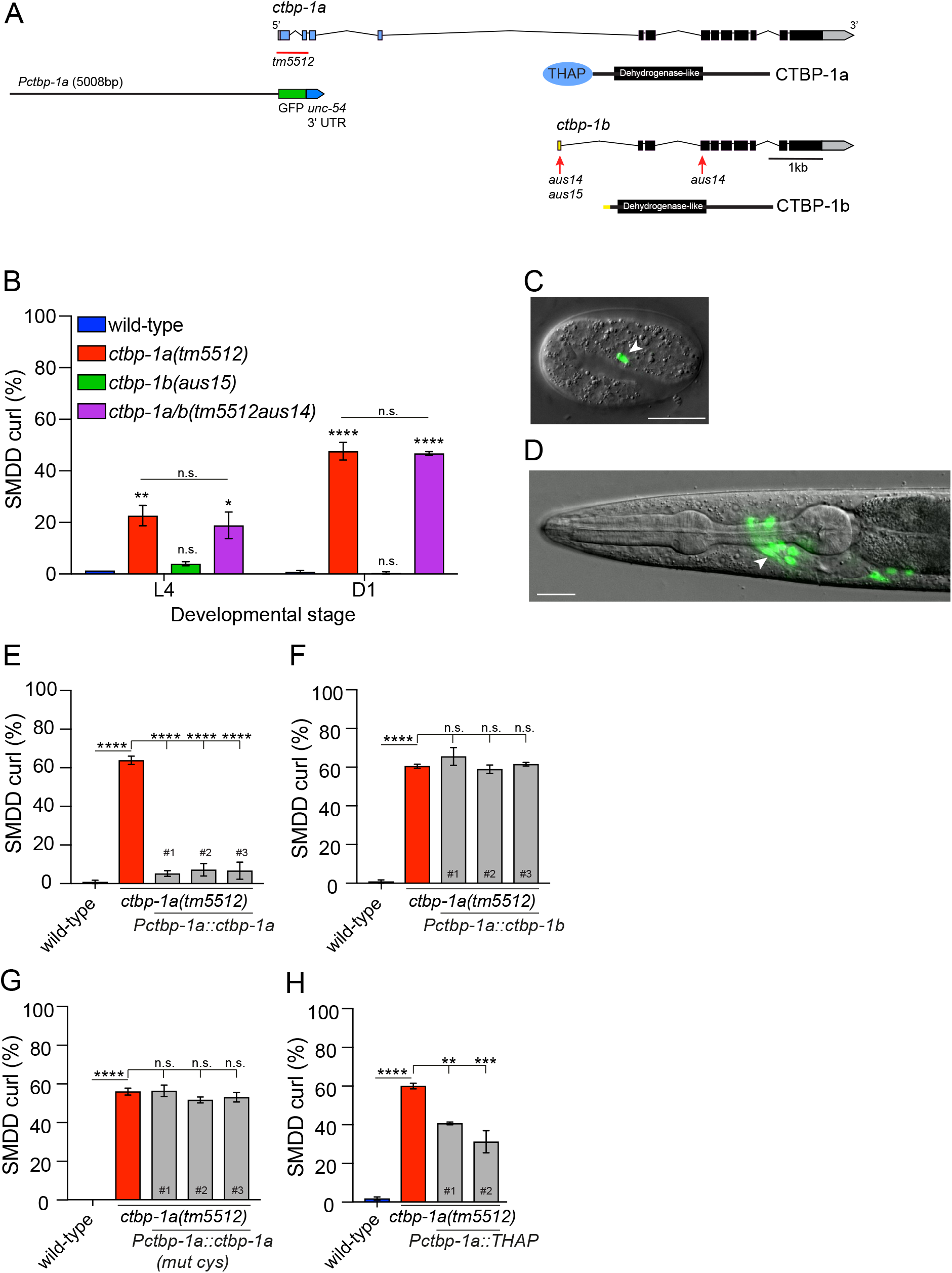
CTBP-1a acts cell-autonomously to control SMDD development. (A) Gene structures and protein domains of *ctbp-1a* and *ctbp-1b*. Shared regions (black), *ctbp-1a*-specific (blue) and *ctbp-1b*-specific (yellow). Genetic lesions used in this study (red bar and arrows). 5008 bp promoter used for *ctbp-1a* expression analysis (black line). (B) Quantification of the SMDD axon curl phenotype in wild-type and *ctbp-1* mutants at larval stage 4 (L4) and adult day 1 (D1). Data presented as mean ± S.E.M (bar) of 3 biological replicates, n>80 axons. *p<0.05, **p<0.01, ****p<0.0001, n.s. – not significant (one-way ANOVA with Tukey’s correction). (C-D) Expression of a *Pctbp-1a::GFP* transcriptional reporter at the bean stage of embryogenesis (C) and L4 larval stage (D). Nomarski and fluorescence images are overlaid. Arrowheads = SMDD neuron. Scale bars, 20μm. (E) Quantification of SMDD axon curl phenotype of *ctbp-1a(tm5512)* rescue: *ctbp-1a* cDNA under the *ctbp-1a* promoter (5008 bp) rescues the *ctbp-1a(tm5512)* SMDD curl phenotype of day 2 adults (3 independent transgenic rescue lines in grey). Data presented as mean ± S.E.M (bar) of 3 biological replicates, n>80 axons. ****p<0.0001 (one-way ANOVA with Tukey’s correction). (F) Quantification of SMDD axon curl phenotype of *ctbp-1a(tm5512)* rescue: *ctbp-1b* cDNA under the *ctbp-1a* promoter does not rescue the *ctbp-1a(tm5512)* SMDD curl phenotype of day 2 adults (3 independent transgenic rescue lines in grey). Data presented as mean ± S.E.M (bar) of 3 biological replicates, n>80 axons. ****p<0.0001, n.s. – not significant (one-way ANOVA with Tukey’s correction). (G) Quantification of SMDD axon curl phenotype of *ctbp-1a(tm5512)* rescue: expression of *ctbp-1a(mut cys)* under the *ctbp-1a* promoter does not rescue the *ctbp-1a(tm5512)* SMDD curl phenotype of day 2 adults (3 independent transgenic rescue lines in grey). Data presented as mean ± S.E.M (bar) of 3 biological replicates, n>80 axons. ****p<0.0001, n.s. – not significant (one-way ANOVA with Tukey’s correction). (H) Quantification of SMDD axon curl phenotype of *ctbp-1a(tm5512)* rescue: expression of *ctbp-1a(THAP)* cDNA under the *ctbp-1a* promoter partially rescues the *ctbp-1a(tm5512)* SMDD curl phenotype (2 independent transgenic rescue lines in grey). Data presented as mean ± S.E.M (bar) of 3 biological replicates, n>80 axons. **p<0.01, ***p<0.001, ****p<0.0001, n.s. – not significant (one-way ANOVA with Tukey’s correction).

### CTBP-1a acts cell-autonomously to control SMDD development

To examine where *ctbp-1a* is expressed, we generated a transcriptional *gfp* reporter using 5008 bp of the *ctbp-1a* promoter *(Pctbp-1a::GFP)*. We found that the *Pctbp-1a::GFP* transgene is first detectable in the SMDD neurons at the bean stage of embryogenesis (Figure 2C). In L4 larvae, the *Pctbp-1a::GFP* transgene drives expression in 12 head neurons, including the SMDD and SMDV neurons (Figure 2D). The presence of SMDV expression prompted us to examine whether CTBP-1a controls axonal development in the ventral SMDs as it does in the dorsal SMDs. Indeed, we found that the SMDV neurons exhibit developmental defects in *ctbp-1a(tm5512)* animals (Figure S2A-B).

The neuronal expression pattern of *Pctbp-1a::GFP* suggests that CTBP-1a regulates SMDD development autonomously. To examine this, we first performed transgenic rescue experiments showing that driving *ctbp-1a* cDNA with the *ctbp-1a* promoter fully rescued SMDD axonal defects of *ctbp-1a(tm5512)* mutant animals (Figure 2E and S2C). Rescue was also observed when driving *ctbp-1a* cDNA using the *lad-2* promoter, which drives expression in the SMDs and 14 other neurons (Aurelio et al., 2002) (Figure S2D). Together, these data suggest that CTBP-1a regulates SMDD development cell-autonomously.

As predicted by our analysis of the *ctbp-1b(aus15)* mutant, we found that driving *ctbp-1b* expression with the *ctbp-1a* promoter does not rescue the SMDD axonal defects of *ctbp-1a(tm5512)* mutant animals (Figure 2F). Because CTBP-1b lacks the THAP DNA binding domain, we examined whether this domain is necessary and sufficient for CTBP-1a regulation of SMDD development. THAP-containing proteins are defined by a zinc-coordinating cysteine-containing consensus module that is important for sequence-specific DNA binding (Clouaire et al., 2005). We mutated two of the cysteines within the full-length CTBP-1a (C5A, C10A) to disrupt THAP-domain function and found that this abrogated the ability of CTBP-1a to rescue the *ctbp-1a(tm5512)* SMDD axonal defect (Figure 2G). Next, we exclusively expressed the THAP domain using the *ctbp-1a* promoter in *ctbp-1a(tm5512)* animals and found that the SMDD axonal defects were partially rescued (Figure 2H). These data support the requirement for the CTBP-1a THAP domain in regulating SMDD development.

CTBP-1a also houses a conserved PXDLS-binding motif, which is important for interactions with CTBP-1-binding proteins (Nardini et al., 2003). We generated an A203E mutation in the PXDLS-binding motif of full-length CTBP-1a, which was previously shown to abrogate interactions with PXDLS-containing binding proteins (Nardini et al., 2003; Nicholas et al., 2008). We found that the A203E mutation has no detectable effect on the rescuing ability of CTBP-1a in the context of SMDD development (Figure S2E). Together, our data suggest that the intrinsic DNA-binding capacity of the CTBP-1a THAP domain, potentially independent of corepressor proteins, is critical for the function of CTBP-1a in controlling SMDD development.

### The L1CAM LAD-2 and CTBP-1a act in parallel to control SMDD development

L1 cell adhesion molecules (L1CAMs) are critical regulators of nervous system development and maintenance. A previous study showed that LAD-2, a *C. elegans* L1CAM ortholog, controls axon guidance of the SMD, SDQL/R and PLN sublateral neurons (Wang et al., 2008). Therefore, we examined whether the functions of CTBP-1a and LAD-2 in controlling SMDD development are related. We found, however, that SMDD length of *lad-2(tm3056)* null mutant animals is similar to wild type at the L4 and 1-day adult stages and that the *lad-2(tm3056); ctbp-1a(tm5512)* double mutant SMDD length is no longer than the *ctbp-1a(tm5512)* single mutant (Figure 3A). Like *ctbp-1a* mutants, *lad-2(tm3056)* animals exhibit abnormal SMD axon trajectories (curl phenotype) (Figure 3B) (Wang et al., 2008). We therefore asked whether *lad-2* and *ctbp-1a* control SMDD axon guidance through the same genetic pathway. We found that *lad-2(tm3056)* and *ctbp-1a(tm5512)* single mutants exhibit similar penetrance of SMDD curl phenotype at the L4 stage (Figure 3B), although the *lad-2(tm3056)* curls occur earlier (L1 stage) than in *ctbp-1a(tm5512)* animals (L4 stage) (Figure 1H) (Wang et al., 2008). Interestingly, loss of *ctbp-1a* but not *lad-2* causes an increase in SMDD defects between the L4 and adult stage, suggesting that they function in separate pathways to control SMDD development (Figure 3B). Confirming this hypothesis, the SMDD defects were additive in the *lad-2(tm3056); ctbp-1a(tm5512)* double mutant when compared to either single mutant (Figure 3B). However, both CTBP-1a and LAD-2 act cell autonomously to control SMDD development, as driving *lad-2* expression with either the *lad-2* or *ctbp-1a* promoter fully rescued the *lad-2(tm3056)* SMDD axon guidance defect (Figure S3A). Taken together, our data show that CTBP-1a and LAD-2/L1CAM act cell-autonomously but in parallel genetic pathways to control SMDD development.

**Figure 3.**
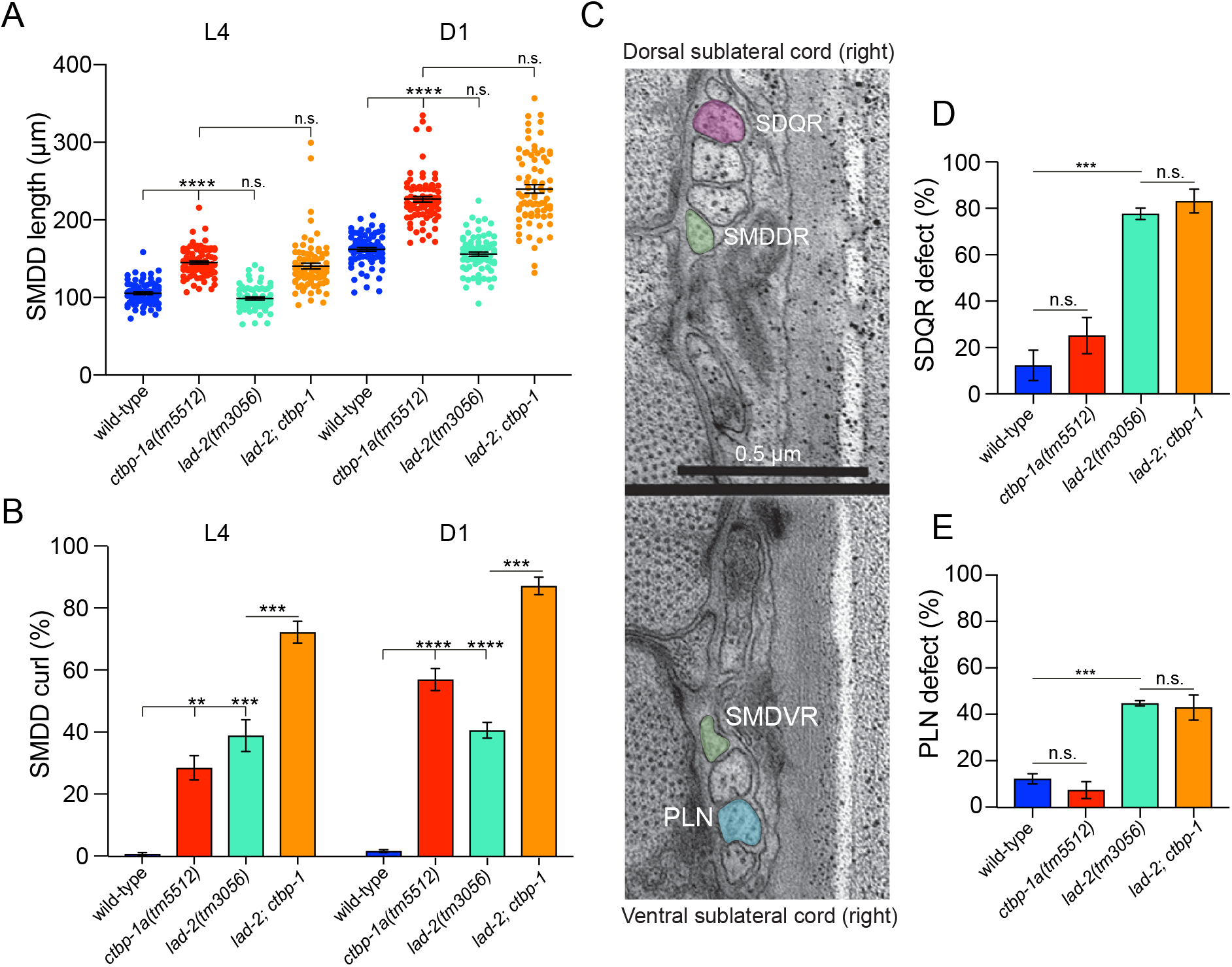
CTBP-1a acts in a parallel genetic pathway to LAD-2/L1CAM. (A) Quantification of SMDD length of wild-type and mutant animals at larval stage 4 (L4) and adult day 1 (D1). Data presented as individual axon lengths (points) with mean ± S.E.M (bar). n=67-82 axons. ****p<0.0001, n.s. – not significant (one-way ANOVA with Tukey’s correction). (B) Quantification of the SMDD curl phenotype of wild-type and mutant animals at larval stage 4 (L4) and adult day 1 (D1). Data presented as mean ± S.E.M (bar) of 3 biological replicates, n>100 axons. **p<0.01, ***p<0.001, ****p<0.0001, n.s. – not significant (one-way ANOVA with Tukey’s correction). (C) SDQR axons (pink) and PLN axons (blue) extend along the right sublateral cords with the SMDD and SMDV axons (green). Scale bar, 0.5μm. (D-E) Quantification of SDQR (D) and PLN (E) defects of day 1 adults. Data presented as mean ± S.E.M (bar) of 3 biological replicates, n>100 axons. ***p<0.001, n.s. – not significant (one-way ANOVA with Tukey’s correction). SDQ and PLN morphology was visualized by a fluorescent reporter – *otEx331[Plad-2::GFP]*.

As mentioned previously, the SMD neurons extend axons adjacent to those of the SDQL/R and PLN neurons within the sublateral cord (Figure 3C) (White et al., 1986). Because LAD-2/L1CAM also controls SDQL/R and PLN axonal development we asked whether CTBP-1a exhibits functional overlap in these neurons (Wang et al., 2008). We found however that the SDQL/R and PLN axons develop normally in *ctbp-1a(tm5512)* mutant animals (Figure 3D-E and S3B). These data indicate that loss of *ctbp-1a* does not generally disrupt axon guidance within the sublateral cord, rather it causes defects specifically in the SMDD neurons.

### CTBP-1 controls SMDD development by repressing SAX-7/L1CAM expression

Due to the important role of LAD-2/L1CAM in SMDD development, we hypothesized that the other *C. elegans* L1CAM ortholog SAX-7 may also function in this regard. However, a previous study showed that unlike LAD-2, SAX-7 is not required for SMD, SDQL/R or PLN development and that loss of *sax-7* does not affect *lad-2* phenotypes in these neurons (Wang et al., 2008). Nevertheless, it is known that inappropriate expression of guidance receptors and their ligands can disrupt neuronal development (Colavita et al., 1998; Hamelin et al., 1993). We therefore hypothesized that CTBP-1, a known transcriptional corepressor, may limit SAX-7 expression to enable faithful SMDD development.

The *sax-7* locus encodes long and short protein isoforms that play multiple roles in axonal maintenance and dendritic branching (Figure 4A) (Benard et al., 2012; Dong et al., 2013; Pocock et al., 2008; Salzberg et al., 2013; Sasakura et al., 2005). The longer SAX-7 isoform (SAX-7L) contains 6 Ig-like domains, 5 FnIII domains and a cytoplasmic tail that houses an ankyrin binding motif, a FERM domain and a PDZ domain (Figure 4A). The shorter SAX-7 isoform (SAX-7S) lacks the first two Ig-like domains (Figure 4A). Using the *sax-7(eq1)* allele, which affects both SAX-7 isoforms, we confirmed that SMDD development is not dependent on SAX-7 (Figure 4A-B). Remarkably, however, loss of *sax-7* partially suppresses the *ctbp-1a(tm5512)* SMDD curl phenotype (Figure 4B). We confirmed suppression of the *ctbp-1a(tm5512)* SMDD curl phenotype using an independently isolated *sax-7(nj48)* allele, which like *eq1*, affects both SAX-7 isoforms (Figure 4A-B). We next asked whether removal of a specific SAX-7 isoform could suppress the *ctbp-1a(tm5512)* SMDD curl phenotype. To this end, we combined the *ctbp-1a(tm5512)* mutation with either the SAX-7L-specific mutation *(nj53)* or the SAX-7S-specific mutation *(ot820)* (Rahe, 2019; Sasakura et al., 2005). We found that removal of SAX-7S but not SAX-7L suppresses the *ctbp-1a(tm5512)* SMDD curl phenotype (Figure 4B). Further, removal of *sax-7* partially rescued the highly penetrant SMDD curl phenotype of *ctbp-1a(tm5512); lad-2(tm3056)* animals (Figure S4A). Inappropriate expression of *sax-7s* in the SMDDs causes the *ctbp-1a(tm5512)* mutant phenotype as low-level expression (0.2ng/μl) of *sax-7s*, under the *ctbp-1a* promoter, restored the SMDD curl phenotype to *ctbp-1a(tm5512); sax-7*(*ot820)* animals, without causing defects in wild-type animals (Figure 4C). We next asked whether the increase of SMDD length in *ctbp-1a(tm5512)* animals is also dependent on SAX-7S. We found, however, that the *sax-7(ot820)* mutation does not reduce the SMDD overgrowth phenotype of the *ctbp-1a(tm5512)* mutant in L4 larvae (Figure S4B). These data suggest that the molecular mechanism(s) through which CTBP-1a controls SMDD outgrowth and guidance are distinct.

**Figure 4.**
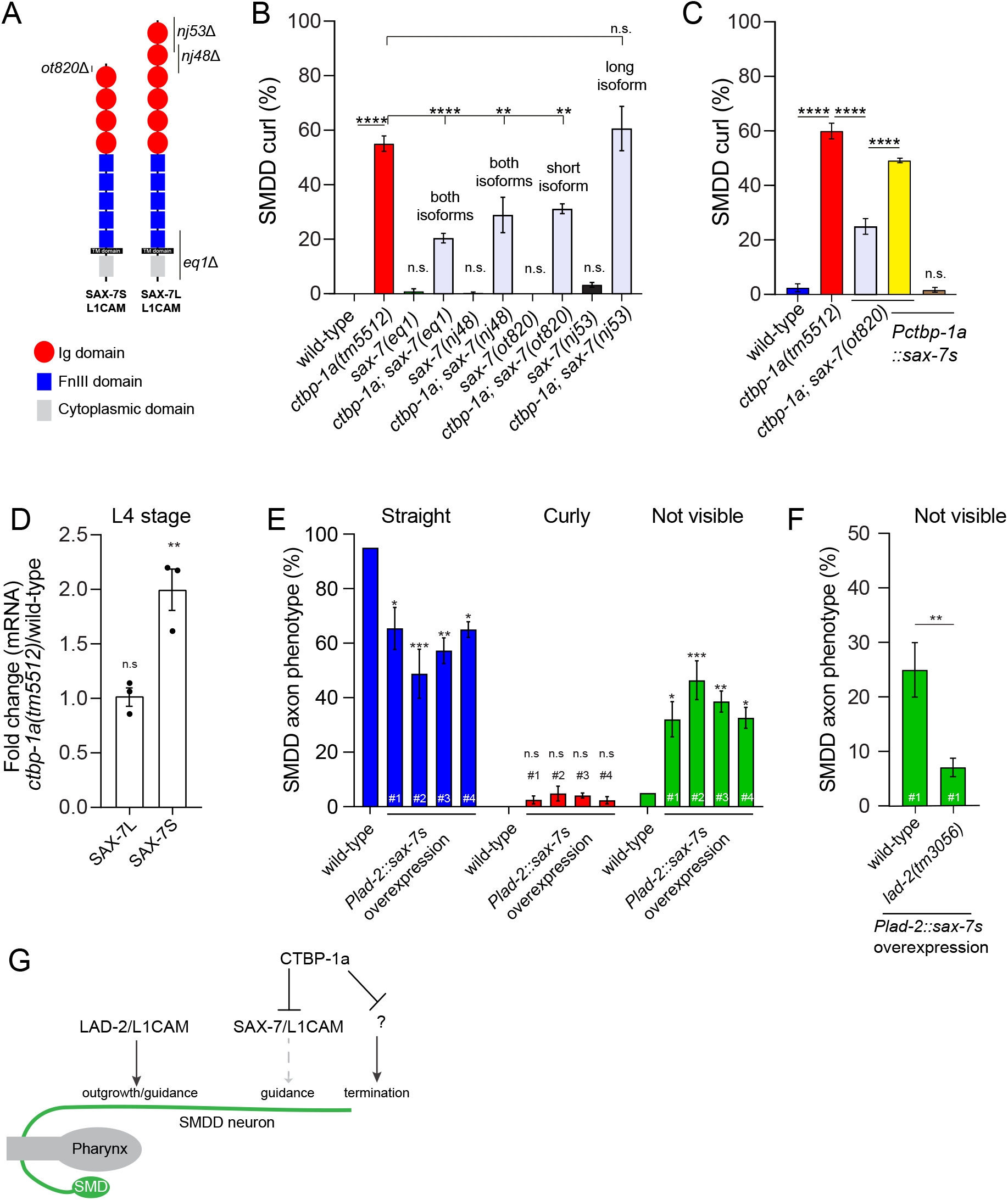
CTBP-1a regulates *sax-7s* for correct SMDD development. (A) Protein structure of SAX-7S and SAX-7L showing the genetic lesions used in this study (black bars). Ig domain, immunoglobulin domain, FnIII domain, fibronectin domain III, cytoplasmic domain (FERM domain, Ankyrin-binding domain, PDZ domain). (B) Quantification of the SMDD axon curl phenotype in *sax-7* single and double mutants at adult day 2 (D2). Data presented as mean ± S.E.M (bar) of 3 biological replicates, n>100 axons. **p<0.01, ****p<0.0001, n.s. – not significant (one-way ANOVA with Tukey’s correction). (C) Quantification of the SMDD axon curl phenotype: low expression of *sax-7* under the *ctbp-1a* promoter *(Pctbp-1a::sax-7s* – 0.2ng/μl) restores the SMDD curl phenotype to *ctbp-1a(tm5512); sax-7(ot820)* animals. Data presented as mean ± S.E.M (bar) of 3 biological replicates, n>100 axons. ****p<0.0001, n.s. – not significant (one-way ANOVA with Tukey’s correction). (D) Expression of *sax-7s* and *sax-7L* transcripts in *ctbp-1a(tm5512)* animals relative to wild-type at the L4 stage. qRT-PCR data presented as 3 biological replicates (points) with mean ± S.E.M (bar). **p<0.01, n.s. – not significant (one-way ANOVA with Dunnett’s multiple comparisons test). (E) Quantification of SMDD axon phenotype of *sax-7s* overexpression in neurons *(Plad-2::sax-7s* – 20ng/μl, 4 independent transgenic lines). Data presented as mean ± S.E.M (bar) of 3 biological replicates, n>50 animals. *p<0.05, **p<0.05, ***p<0.001, n.s. – not significant (one-way ANOVA with Dunnett’s correction for each phenotype). (F) Quantification of SMDD not visible axon phenotype of wild-type and *lad-2(tm3056)* animals expressing *Plad-2::sax-7s* overexpression line #1. Data presented as mean ± S.E.M (bar) of 3 biological replicates, n>50 animals. **p<0.05 (unpaired t-test). (G) CTBP-1a represses SAX-7/L1CAM expression to allow correct axon guidance and outgrowth. CTBP-1a regulates SMDD axon termination through an unknown mechanism. In a parallel genetic pathway, LAD-2/L1CAM controls SMDD axon guidance.

Our collective genetic data suggest that the *ctbp-1a(tm5512)* mutant SMDD curl phenotype is caused by elevated *sax-7s* (Figure 4B-C). To examine whether CTBP-1a regulates *sax-7s* expression, we performed quantitative real-time PCR (qRT-PCR) on RNA samples extracted from wild-type and *ctbp-1a(tm5512)* mutant synchronized L4 larvae (Figure 4D). Consistent with our phenotypic data, we found that the level of *sax-7s* but not *sax-7L* is increased in the *ctbp-1a(tm5512)* mutant compared to wildtype (Figure 4D). These data suggest that CTBP-1a represses *sax-7s* expression, directly or indirectly, to enable correct SMDD axon development. If this was the case, one would predict that inappropriate overexpression of SAX-7S in wild-type animals would cause SMDD axon defects. We therefore overexpressed *sax-7s* in wild-type animals and monitored SMDD development. We found that overexpression of *sax-7s* under the *lad-2* promoter caused a severe neomorphic axon outgrowth defect where SMDD axons did not enter the sublateral cord (not visible phenotype), suggesting that the axons did not exit the nerve ring (Figure 4E). We confirmed that overexpressing *sax-7s* did not cause cell death as SMDD cell bodies are present in L4 larvae of *Plad-2::sax-7s* overexpression animals (Figure S5A). In contrast to overexpression in neurons, overexpression of *sax-7s* in the hypodermis *(dpy-7* promoter) or body wall muscle *(myo-3* promoter), other tissues in which *sax-7* is expressed, had no detectable effect on SMDD development (Figure S5B-C) (Chen et al., 2001). The neomorphic axon outgrowth phenotype caused by *sax-7s* overexpression in the nervous system was not previously observed in animals lacking *ctbp-1, lad-2* or *sax-7*. However, we hypothesized that the axon outgrowth phenotype caused by *sax-7s* overexpression in the SMDD neurons may be due to an inappropriate *in cis* interaction between the two *C. elegans* L1CAMs. To test this hypothesis, we crossed the *lad-2(tm3056)* mutation into one of the *sax-7s* overexpression lines (Figure 4F). Removing *lad-2* from animals overexpressing *Plad-2::sax-7s* significantly increased the number of SMDDs axons entering the sublateral cord (Figure 4F). These data show that the neomorphic effect caused by *sax-7s* overexpression is dependent on LAD-2. Together, these data reveal that CTBP-1a repression of the short isoform of SAX-7/L1CAM enables LAD-2/L1CAM-driven SMDD axon development.

## DISCUSSION

In this study, we found that axons of the *C. elegans* sublateral SMDD motor neurons terminate outgrowth at the first day of adulthood, after which the axons partially retract. Animals lacking the CTBP-1 transcriptional corepressor have longer SMDD axons than wild-type that fail to retract in adults. Loss of CTBP-1 also causes progressive misguidance of SMDD axons outside the sublateral tract. We found that CTBP-1 controls SMDD development by repressing expression of the short isoform of SAX-7/L1CAM. Repression of SAX-7 is critical for SMDD development as its overexpression causes severe defects in SMDD outgrowth that are genetically dependent on the other *C. elegans* L1CAM, LAD-2. Further, we found that LAD-2 acts in a parallel pathway to CTBP-1 and SAX-7/L1CAM to control SMDD development.

Hence, the *presence* of LAD-2/L1CAM and *absence* of SAX-7/L1CAM is required for faithful SMDD development (Figure 4G).

### Developmental Plasticity of the SMDD Axons

Our study reveals that the SMDD axons continuously extend during larval development and into the first day of adulthood. We further found that SMDD axon extension does not directly scale with worm body length, suggesting cell-autonomous control of SMDD length. Why the SMDD axons subsequently retract from day 1 to day 3 of adulthood is unknown. Perhaps the SMDDs perform differential functions in early and late adult life such that their axons need to be located in distinct environments or that they need to modify synaptic connectivity during this period. The ability of the SMDD axons to remodel may also point to a role in experience-dependent learning in response to changing environments. The reported roles for the SMDD neurons in coordinating behavior and circuit function support this hypothesis (Gray et al., 2005; Shen et al., 2016; Yeon et al., 2018).

### Parallel Regulation of Axon Development by Two L1CAMs

Our work shows that CTBP-1a regulates expression of the short isoform of SAX-7/L1CAM. We found that removal of the SAX-7S but not SAX-7L suppresses the SMDD curl phenotype of *ctbp-1a* mutant animals. Correct regulation of SAX-7S expression is critical, as inappropriate expression of SAX-7S in the SMDDs causes severe axon outgrowth defects such that they do not exit the nerve ring. In contrast, overexpressing the SAX-7S in neighboring hypodermis and body wall muscle, which the SMDDs potentially use as a growth substrate (White et al., 1986), has no visible effect on SMDD development. These data suggest that under standard laboratory conditions, preventing expression of SAX-7S/L1CAM in the SMDDs is required for their development.

Our data show that CTBP-1a regulation of SMDD development through SAX-7S occurs within a distinct temporal window to LAD-2 – the other *C. elegans* L1CAM. SMDD axon guidance defects caused by loss of LAD-2/L1CAM appear in early L1 larvae (Wang et al., 2008), whereas the effects of CTBP-1a loss are not detected until the L4 stage. This suggests that correct development of the SMDD neurons requires precise control of both *C. elegans* L1CAMs over different timescales. Our experimental evidence further supports this hypothesis: 1) using the *lad-2* promoter to overexpress *sax-7s* causes severe SMDD outgrowth defects, further supporting a requirement for repression of *sax-7s* expression in *lad-2*-expressing neurons; 2) SMDD axon defects caused by *sax-7s* overexpression are dependent on *lad-2*. These data suggest that expression of SAX-7S and LAD-2 within the same neuron causes inappropriate adhesion and/or signaling that severely affects axon outgrowth. Could CTBP-1a repression of SAX-7S in the SMDD neurons also provide functional/structural flexibility? Perhaps under certain environmental or stress states expression of SAX-7S could be advantageous by providing post-developmental structural integrity to the SMDD neurons or by stimulating axon retraction. For such a scenario, reduction of CTBP-1a levels would promote expression of SAX-7S in the SMDDs. Indeed, CtBP in mice undergoes proteasome-dependent degradation under stress, which may provide alternative strategies for neuronal survival or signaling (Zhang et al., 2003).

### Regulation of L1CAMs by THAP-containing Proteins is Potentially Conserved

The CTBP-1a protein contains an N-terminal THAP domain that is defined by a C2CH zinc-dependent DNA binding motif. THAP domain-containing proteins can act as transcriptional repressors or corepressors either by directly binding to DNA through the C2CH motif or by recruiting corepressor proteins (Clouaire et al., 2005). Our rescue experiments show that mutation of cysteine residues in the C2CH THAP motif of CTBP-1a inhibits its ability to rescue *ctbp-1a(tm5512)* SMDD defects. Since mutation of the C2CH motif abrogates the DNA-binding activity of THAP proteins, this observation suggests that CTBP-1a can directly regulate transcription (Clouaire et al., 2005). Importantly, these cysteine mutations do not detectably cause THAP1 protein instability (Clouaire et al., 2005). We further found that expressing the CTBP-1a THAP domain is alone sufficient to partially rescue SMDD defects of *ctbp-1a(tm5512)* animals. Cumulatively, these data suggest that the THAP domain of CTBP-1a can directly coordinate transcriptional repression, and potentially directly repress gene expression in the SMDDs.

Studies in mammalian models imply that the function of THAP domains in controlling nervous system development and regulation of L1CAM-related molecules may be conserved. Mutations in the THAP1 gene are associated with dystonia, a brain disorder characterized by involuntary muscle contraction (Zakirova et al., 2018). Additionally, mammalian models of THAP1 loss-of-function reveal defects in motor function and anxiety-related behavior (Frederick et al., 2019; Ruiz et al., 2015). In these genetic models, THAP1 heterozygosity can cause a decrease in neuron number within the dentate nucleus of the cerebellum (Ruiz et al., 2015). Additionally, *in vitro* analysis of THAP1 heterozygous striatal neurons show that neurite growth is defective (Zakirova et al., 2018). These data suggest a common function for THAP proteins in the control of nervous system development and behavior. Additional genomic analysis shows that the regulatory mechanism we elucidated in *C. elegans* may also be conserved in mammals. Differential gene expression analysis of multiple mouse models reveals that heterozygous loss of THAP1 causes dysregulation of L1CAM and related cell adhesion molecules, including NCAM (Aguilo et al., 2017; Frederick et al., 2019). These transcriptomic data need to be validated. However, ChIP-sequencing data (ModENCODE) also show that THAP1 interacts with the L1CAM locus and therefore may directly regulate expression of this SAX-7 ortholog in mammals (Consortium, 1998; Davis et al., 2018).

Collectively, our results reveal that the CTBP-1a transcriptional corepressor is required for axonal extension of the SMDD neurons. Instead of terminating their outgrowth as animals reach adulthood, *ctbp-1a* mutant SMDD axons continue to extend. In addition to regulating axon outgrowth, CTBP-1a is further required for SMDD guidance within the sublateral nerve cord. We found that CTBP-1a controls SMDD guidance by repressing SAX-7/L1CAM expression. This regulatory relationship is crucial, as inappropriate SAX-7/L1CAM expression causes severe defects in SMDD development that are dependent on the presence of LAD-2/L1CAM. Further, the expression of LAD-2/L1CAM is required for the early stages of SMDD development in parallel to the CTBP-1a/SAX-7 regulatory axis. Taken together, our results show that the expression of L1CAM family members is tightly regulated to shape axon outgrowth and guidance decisions – control mechanisms we have shown to be mediated by CTBP-1a, a THAP domain transcriptional corepressor protein.

## CONTACT FOR REAGENT AND RESOURCE SHARING

Further information and requests for resources and reagents should be directed to and will be fulfilled by the Lead Contact, Roger Pocock.

## EXPERIMENTAL MODEL AND SUBJECT DETAILS

### Mutant and transgenic reporter strains

Strains were grown using standard growth conditions on NGM agar at 20°C or 25°C on *Escherichia coli* OP50 (Sulston and Brenner, 1974). Neuroanatomical reporter strains used – *rhIs4 Is[Pglr-1::GFP], rpEx1739 Ex[Pctbp-1a::GFP], otEx331 Ex[Plad-2::GFP]*. Detailed strain information is available in Table S3. Developmental stage of animals used for each experiment are specified in figure legends.

### Transgenic lines

Rescue constructs were injected into *rhIs4; ctbp-1a(tm5512)* or *rhIs4; lad-2(tm3056)* mutant backgrounds at 2 ng/μl with *Punc-122::GFP* (20 ng/μl) as injection marker. Overexpression constructs were injected into *rhIs4 (Pglr-1::GFP)* background at 5-20 ng/μl with *Punc-122::GFP* (20 ng/μl) as injection marker. The *Pctbp-1a::GFP* expression construct was injected into N2 (wild-type) background at 50 ng/μl with *Pttx-3::dsRed2* (50 ng/μl) as injection marker. Microinjections were performed using standard methods (Mello et al., 1991). Briefly, young adult worms were picked to an agarose pad covered with oil on a glass slide. The immobilized worms were injected using FemtoJet 4x injector (Eppendorf) controlled by InjectMan 4 (Eppendorf). Detailed strain information is available in Table S3.

## METHOD DETAILS

### Molecular cloning

All cloning and mutagenesis was performed using In-Fusion^®^ restriction-free cloning (Takara). Linear PCR products and/or vectors (as detailed below) were fused using restriction-free In-Fusion^®^ HD cloning reagents. Plasmid sequences were confirmed using Sanger sequencing.

#### RJP383 *Pctbp-1a::GFP*

The *Pctbp-1a::GFP* reporter construct was generated by cloning the 5008 bp *ctbp-1a* promoter from worm genomic DNA into the promoter-less GFP pPD95.75 expression vector (linearised by HindIII and XbaI).

#### RJP414 *PCTBP-1a::sax-7s cDNA*

RJP414 was generated by amplifying the pRP13 *Pdpy-7::sax-7* vector minus the *dpy-7* promoter sequence and amplifying the 5008 bp *ctbp-1a* promoter from RJP383.

#### RJP422 *Pctbp-1a::ctbp-1a::mCherry*

First, the RJP420 *ctbp-1a::mCherry* vector was generated by amplifying the pPD95.75 mCherry expression vector and amplifying the 2181 bp *ctbp-1a* cDNA from pAER019 *Pglr-1::ctbp-1a::V5::ctbp-1* 3 UTR (Reid et al., 2015). RJP422 was then generated by cloning the 5008 bp *ctbp-1a* promoter from RJP383 into the RJP420 vector (linearised by BamHI).

#### RJP423 *Pctbp-1a::ctbp-1b::mCherry*

First, the RJP421 *ctbp-1b::mCherry* vector was generated by amplifying the pPD95.75 mCherry expression vector and amplifying the 1818 bp *ctbp-1b* cDNA from worm cDNA. RJP422 was then generated by cloning the 5008 bp *ctbp-1a* promoter from RJP383 into RJP421 vector (linearized by BamHI).

#### RJP424 *Plad-2::ctbp-1a::mCherry*

RJP424 was generated by cloning the 4063 bp *lad-2* promoter from worm genomic DNA into the RJP420 vector (linearized with BamHI).

#### RJP424 *Pdpy-7::ctbp-1a::mCherry*

RJP424 was generated by cloning the 249 bp *dpy-7* promoter from pTB80 *Pdpy-7::GFP* into the RJP420 vector (linearised by BamHI).

#### RJP515 *Plad-2::sax-7s*

RJP515 was generated by amplifying the *sax-7s* cDNA sequence from RJP414 and amplifying the *lad-2* promoter from RJP424.

#### RJP514 *Pctbp-1a::ctbp-1a(A203E)::mCherry*

RJP514 was generated using site-directed mutagenesis to mutate the key alanine residue to glutamic acid in the PXDLS-binding cleft domain in the *Pctbp-1a::ctbp-1a cDNA::mCherry* vector (Nicholas et al., 2008).

#### RJP426 *Pctbp-1a::ctbp-1a(THAP only)::mCherry*

RJP426 was generated by amplifying the sequence corresponding to the 140 amino acid THAP domain of the *Pctbp-1a::ctbp-1a cDNA::mCherry* vector (minus the *ctbp-1b*-shared sequence).

#### RJP427 *Pctbp-1a::ctbp-1a(C5A,C10A)::mCherry*

RJP427 was generated using site-directed mutagenesis to mutate key cysteines at position 5 and 10 to alanine in the CTBP-1a THAP domain in the *Pctbp-1a::ctbp-1a cDNA::mCherry* vector (Clouaire et al., 2005).

#### RJP540 *Pmyo-3::sax-7s*

RJP540 was generated by amplifying the *myo-3* promoter sequence from pPD95.86-*myo-3* and amplifying the *sax-7s* cDNA from RJP414.

#### RJP517 *Pctbp-1a::lad-2*

RJP517 was generated by amplifying the *Pctbp-1a::GFP* vector (minus GFP) and *lad-2* cDNA from worm cDNA.

#### RJP520 *Plad-2::lad-2*

RJP520 was generated by amplifying the *Pctbp-1a::lad-2 cDNA* vector (minus *Pctbp-1)* and *Plad-2* from worm genomic DNA.

### CRISPR-Cas9

sgRNA target sequences were designed and incorporated into a *pU6::klp-12* sgRNA expression vector by PCR as previously described (Norris et al., 2015). Wild-type (N2) animals were injected with a mix consisting of the sgRNA expression vector(s) (125 ng/μl), Cas9 expression vector *(Peft-3::cas9::tbb-2)* (50 ng/μl), pCFJ90 *(Pmyo-2::mCherry::unc-54)* (2.5 ng/μl) and pCFJ104 *(Pmyo-3::mCherry::unc-54)* (5 ng/μl). PCR and Sanger sequencing was performed on mCherry-expressing animals to identify deletions. Genome modifications generated in this study: *aus14* is a 4 bp deletion in *ctbp-1b* exon 1; *aus15* is a 39 bp deletion in *ctbp-1b* exon 1/intron 1 and 5 bp deletion in *ctbp-1b* exon 4b.

### Microscopy

Animals were anesthetized with 0.2% levamisole hydrochloride on 5% agarose pads, and images were obtained with an Axio Imager M2 fluorescence microscope, Axiocam 506 mono camera and Zen software (Zeiss).

### Phenotypic analyses

#### SMDD axon morphology assays

SMD morphology was analyzed using the *rhIs4 Is[Pglr-1::GFP]* or *rpEx1739 Ex[Pctbp-1a::GFP]* reporters using the 40x objective. SMDD curl (%) indicates the percentage of SMDD axons that ‘curl’ away from or leave the dorsal sublateral path along which the SMDD axons extend. SMDD axonal phenotype (%) measures the proportion of axons exhibiting the three possible phenotypes: ‘Straight’, ‘Curly’ and ‘Not visible’. ‘Straight’ SMDD axons extend along the dorsal sublateral cord; ‘Curly’ SMDD axons leave the dorsal sublateral cord, and ‘Not visible’ means that there is no axon visible at any position along the dorsal sublateral cord. For each genotype, the total percentages of ‘Straight’, ‘Curly’ and ‘Not visible’ phenotypes add up to 100%. For all SMDD assays, 3 biological replicates were performed, and statistical significance was assessed by student’s t-test or one-way ANOVA followed by Tukey’s or Dunnett’s multiple comparisons tests.

#### SDQL/R and PLN axon morphology assays

SDQR, SDQL and PLN axon guidance assays were performed as previously described, using *otEx331 Ex[Plad-2::GFP]* (Wang et al., 2008). The SDQR axon was scored as defective if the axon extended ventrally. The SDQL axon was scored as defective if the axon extended ventrally. The PLN axon was scored as defective if the axon extended posteriorly. 3 biological replicates were performed, and statistical significance was assessed by a one-way ANOVA followed by Tukey’s multiple comparisons tests.

#### SMDD axon length

SMDD axon length images were obtained with a 40x objective using GFP and DIC channels. SMDD axon length (μm) was quantified in FIJI (ImageJ) by tracing from the anterior bulb of the pharynx (DIC images) to the distal tip of the axon (GFP fluorescence images) in DIC/GFP composite images. Two biological replicates were performed and the measurements pooled for analysis. Statistical significance was assessed by student’s t-test or one-way ANOVA followed by Tukey’s multiple comparisons test.

#### Body length

Body length images were obtained with a 20x objective using DIC channel. Body length images were taken at specified developmental stages at 20x magnification. Body length (μm) was quantified in FIJI (ImageJ) by tracing along the middle of the animal from the anterior tip of the head to the tail. When an animal length spanned more than one image, overlapping images were taken and joined together in Adobe Photoshop. Two biological replicates were performed and the measurements pooled for analysis. Statistical significance was assessed by student’s t-test for each developmental stage.

### qRT-PCR assays

Total RNA of L4 stage worms was isolated using the RNAeasy mini kit (Qiagen 74104), according to manufacturer’s instructions. Total cDNA was obtained using oligodT primers and the ImProm-II™ Reverse Transcription System (A3800) followed by quantitative PCR using SYBR green (Thermo Scientific 4385610) and Light Cycler 480 (Roche). Samples from three biological replicates were run in triplicate. The *C. elegans* reference gene *pmp-3* was used as an internal control. Primer sequences are listed in the Key Resources Table.

## QUANTIFICATION AND STATISTICAL ANALYSIS

All experiments were performed in three independent replicates, unless specified. The numbers of animals analyzed for specific experiments are reported in the figures or legends. Statistical analysis was performed in GraphPad Prism 8 using unpaired student’s t-test, or one-way analysis of variance (ANOVA) for comparison followed by Tukey’s Multiple Comparison Test or Dunnett’s Multiple Comparison Test, where applicable. Values are expressed as mean ± S.E.M. Differences with a p value <0.05 were considered significant.

## AUTHOR CONTRIBUTIONS

T.S. conducted the experiments; T.S., H.R.N and R.P. designed the experiments and wrote the paper.

## ACKNOWLEDGEMENTS

We thank members of the Pocock Laboratory for comments on the manuscript. We thank David Hall for extraction and annotation of electron micrographs. Some strains were provided by the Caenorhabditis Genetics Center (University of Minnesota), which is funded by NIH Office of Research Infrastructure Programs (P40 OD010440). This research was supported by an Australian Government Research Training Program (RTP) Scholarship to T.S. This work was supported by the following grants: NHMRC (Project GNT1105374 and Senior Research Fellowship GNT1137645 to R.P.) and veski Innovation Fellowship (VIF23 to R.P.).

## REFERENCES

Aguilo, F., Zakirova, Z., Nolan, K., Wagner, R., Sharma, R., Hogan, M., Wei, C. G., Sun, Y. F., Walsh, M. J., Kelley, K., et al. (2017). THAP1: Role in Mouse Embryonic Stem Cell Survival and Differentiation. Stem Cell Rep 9, 92–107.

Aurelio, O., Hall, D. H. and Hobert, O. (2002). Immunoglobulin-domain proteins required for maintenance of ventral nerve cord organization. Science 295, 686–690.

Bagri, A., Cheng, H. J., Yaron, A., Pleasure, S. J. and Tessier-Lavigne, M. (2003). Stereotyped pruning of long hippocampal axon branches triggered by retraction inducers of the semaphorin family. Cell 113, 285–299.

Benard, C. Y., Blanchette, C., Recio, J. and Hobert, O. (2012). The secreted immunoglobulin domain proteins ZIG-5 and ZIG-8 cooperate with L1CAM/SAX-7 to maintain nervous system integrity. PLoS genetics 8, e1002819.

Brummendorf, T., Kenwrick, S. and Rathjen, F. G. (1998). Neural cell recognition molecule L1: from cell biology to human hereditary brain malformations. Curr Opin Neurobiol 8, 87–97.

Chen, L., Ong, B. and Bennett, V. (2001). LAD-1, the Caenorhabditis elegans L1CAM homologue, participates in embryonic and gonadal morphogenesis and is a substrate for fibroblast growth factor receptor pathway-dependent phosphotyrosine-based signaling. J Cell Biol 154, 841–856.

Clouaire, T., Roussigne, M., Ecochard, V., Mathe, C., Amalric, F. and Girard, J. P. (2005). The THAP domain of THAP1 is a large C2CH module with zinc-dependent sequencespecific DNA-binding activity. Proc Natl Acad Sci U S A 102, 6907–6912.

Cohen, N. R., Taylor, J. S., Scott, L. B., Guillery, R. W., Soriano, P. and Furley, A. J. (1998). Errors in corticospinal axon guidance in mice lacking the neural cell adhesion molecule L1. Curr Biol 8, 26–33.

Colavita, A., Krishna, S., Zheng, H., Padgett, R. W. and Culotti, J. G. (1998). Pioneer axon guidance by UNC-129, a C. elegans TGF-beta. Science 281, 706–709.

Consortium, C. e. S. (1998). Genome sequence of the nematode C. elegans: a platform for investigating biology. Science 282, 2012–2018.

Cook, S. J., Jarrell, T. A., Brittin, C. A., Wang, Y., Bloniarz, A. E., Yakovlev, M. A., Nguyen, K. C. Q., Tang, L. T., Bayer, E. A., Duerr, J. S., et al. (2019). Whole-animal connectomes of both Caenorhabditis elegans sexes. Nature 571, 63–71.

Davis, C. A., Hitz, B. C., Sloan, C. A., Chan, E. T., Davidson, J. M., Gabdank, I., Hilton, J. A., Jain, K., Baymuradov, U. K., Narayanan, A. K., et al. (2018). The Encyclopedia of DNA elements (ENCODE): data portal update. Nucleic acids research 46, D794–D801.

Dong, X., Liu, O. W., Howell, A. S. and Shen, K. (2013). An extracellular adhesion molecule complex patterns dendritic branching and morphogenesis. Cell 155, 296–307.

Frederick, N. M., Shah, P. V., Didonna, A., Langley, M. R., Kanthasamy, A. G. and Opal, P. (2019). Loss of the dystonia gene Thap1 leads to transcriptional deficits that converge on common pathogenic pathways in dystonic syndromes. Human molecular genetics 28, 1343–1356.

Gray, J. M., Hill, J. J. and Bargmann, C. I. (2005). A circuit for navigation in Caenorhabditis elegans. Proc Natl Acad Sci U S A 102, 3184–3191.

Hamelin, M., Zhou, Y. W., Su, M. W., Scott, I. M. and Culotti, J. G. (1993). Expression of the Unc-5 Guidance Receptor in the Touch Neurons of C-Elegans Steers Their Axons Dorsally. Nature 364, 327–330.

Hutter, H. (2019). Formation of longitudinal axon pathways in Caenorhabditis elegans. Semin Cell Dev Biol 85, 60–70.

Luo, L. and O’Leary, D. D. (2005). Axon retraction and degeneration in development and disease. Annu Rev Neurosci 28, 127–156.

Mello, C. C., Kramer, J. M., Stinchcomb, D. and Ambros, V. (1991). Efficient gene transfer in C.elegans: extrachromosomal maintenance and integration of transforming sequences. Embo J 10, 3959–3970.

Nagaraj, K., Mualla, R. and Hortsch, M. (2014). The L1 family of cell adhesion molecules: a sickening number of mutations and protein functions. Adv Neurobiol 8, 195–229.

Nardini, M., Spano, S., Cericola, C., Pesce, A., Massaro, A., Millo, E., Luini, A., Corda, D. and Bolognesi, M. (2003). CtBP/BARS: a dual-function protein involved in transcription co-repression and Golgi membrane fission. The EMBO journal 22, 3122–3130.

Nicholas, H. R., Lowry, J. A., Wu, T. and Crossley, M. (2008). The Caenorhabditis elegans protein CTBP-1 defines a new group of THAP domain-containing CtBP corepressors. Journal of molecular biology 375, 1–11.

Norris, A. D., Kim, H. M., Colaiacovo, M. P. and Calarco, J. A. (2015). Efficient Genome Editing in Caenorhabditis elegans with a Toolkit of Dual-Marker Selection Cassettes. Genetics 201, 449–458.

Pocock, R., Benard, C. Y., Shapiro, L. and Hobert, O. (2008). Functional dissection of the C. elegans cell adhesion molecule SAX-7, a homologue of human L1. Mol Cell Neurosci 37, 56–68.

Rahe, D., Carrera, I., Cosmanescu, F., & Hobert, O. (2019). An isoform-specific allele of the *sax-7* locus. microPublication Biology.

Rapti, G., Li, C., Shan, A., Lu, Y. and Shaham, S. (2017). Glia initiate brain assembly through noncanonical Chimaerin-Furin axon guidance in C. elegans. Nature neuroscience 20, 1350–1360.

Reid, A., Sherry, T. J., Yucel, D., Llamosas, E. and Nicholas, H. R. (2015). The C-terminal binding protein (CTBP-1) regulates dorsal SMD axonal morphology in Caenorhabditis elegans. Neuroscience 311, 216–230.

Ruiz, M., Perez-Garcia, G., Ortiz-Virumbrales, M., Meneret, A., Morant, A., Kottwitz, J., Fuchs, T., Bonet, J., Gonzalez-Alegre, P., Hof, P. R., et al. (2015). Abnormalities of motor function, transcription and cerebellar structure in mouse models of THAP1 dystonia. Human molecular genetics 24, 7159–7170.

Salzberg, Y., Diaz-Balzac, C. A., Ramirez-Suarez, N. J., Attreed, M., Tecle, E., Desbois, M., Kaprielian, Z. and Bulow, H. E. (2013). Skin-derived cues control arborization of sensory dendrites in Caenorhabditis elegans. Cell 155, 308–320.

Sasakura, H., Inada, H., Kuhara, A., Fusaoka, E., Takemoto, D., Takeuchi, K. and Mori, I. (2005). Maintenance of neuronal positions in organized ganglia by SAX-7, a Caenorhabditis elegans homologue of L1. Embo J 24, 1477–1488.

Shen, Y., Wen, Q., Liu, H., Zhong, C., Qin, Y., Harris, G., Kawano, T., Wu, M., Xu, T., Samuel, A. D., et al. (2016). An extrasynaptic GABAergic signal modulates a pattern of forward movement in Caenorhabditis elegans. eLife 5.

Sulston, J. E. and Brenner, S. (1974). The DNA of Caenorhabditis elegans. Genetics 77, 95–104.

Tessier-Lavigne, M. and Goodman, C. S. (1996). The molecular biology of axon guidance. Science 274, 1123–1133.

Wang, X., Zhang, W., Cheever, T., Schwarz, V., Opperman, K., Hutter, H., Koepp, D. and Chen, L. (2008). The C. elegans L1CAM homologue LAD-2 functions as a coreceptor in MAB-20/Sema2 mediated axon guidance. J Cell Biol 180, 233–246.

White, J. G., Southgate, E., Thomson, J. N. and Brenner, S. (1986). The structure of the nervous system of the nematode Caenorhabditis elegans. Philosophical transactions of the Royal Society of London. Series B, Biological sciences 314, 1–340.

Xu, N. J. and Henkemeyer, M. (2009). Ephrin-B3 reverse signaling through Grb4 and cytoskeletal regulators mediates axon pruning. Nature neuroscience 12, 268–276.

Yeon, J., Kim, J., Kim, D. Y., Kim, H., Kim, J., Du, E. J., Kang, K., Lim, H. H., Moon, D. and Kim, K. (2018). A sensory-motor neuron type mediates proprioceptive coordination of steering in C. elegans via two TRPC channels. PLoS biology 16, e2004929.

Zakirova, Z., Fanutza, T., Bonet, J., Readhead, B., Zhang, W., Yi, Z., Beauvais, G., Zwaka, T. P., Ozelius, L. J., Blitzer, R. D., et al. (2018). Mutations in THAP1/DYT6 reveal that diverse dystonia genes disrupt similar neuronal pathways and functions. PLoS genetics 14, e1007169.

Zhang, Q., Yoshimatsu, Y., Hildebrand, J., Frisch, S. M. and Goodman, R. H. (2003). Homeodomain interacting protein kinase 2 promotes apoptosis by downregulating the transcriptional corepressor CtBP. Cell 115, 177–186.

